# Limiting-stress-elimination hypothesis: approach to increase savanna cowpea productivity by stress reduction

**DOI:** 10.1101/594754

**Authors:** Acheampong Atta-Boateng, Graeme P. Berlyn

**Affiliations:** School of Forestry and Environmental Studies, Yale University, New Haven, CT 06511, USA

**Keywords:** Sprengel-Liebig law, non-hormonal biostimulant, fertilizer, osmotic stress

## Abstract

We propose and test the Limiting-Stress-Elimination Hypothesis (LSEH), as a decision axiom to guide in determining the optimal intervention strategy towards yield allocation in a savanna legume. We considered osmotic stress and soil nitrogen (N) limitation, which both characterize Guinea savanna agro-ecozone. We hypothesized that biomass allocation will increase when the limiting stress is eliminated at least until the next limiting stress impacts yield. We assessed responses of *Vigna unguiculata* (L.) Walp (cowpea) to osmotic stress treatments (non-hormonal biostimulant and exogenous metabolite) and N input by leaf level physiology, N-fixing capacity and biomass. The relative increase in biomass (%), and pod yields reveal that osmotic stress (45%) more than nitrogen (13%) is limiting to cowpea growth under Guinea savanna conditions, although N fertilization increased nodulation and maximized PSII quantum yield. In the Sprengel-Liebig’s decrement from the maximum concept, the decrement from the maximum for each stressor must be minimized in order to produce the absolute maximum production. However, this may not be economically feasible in many situations. Conversely, LSEH demonstrates that significant productivity is attainable by eliminating a relatively more limiting stress, osmotic stress, regardless of the limitation and natural demand for the relatively less limiting N in leguminous cowpea in the savanna.

## 1. Introduction

Sustainably overcoming food insecurity amid climate threats and rising global population require careful strategic interventions in crop production (Tilman et al., 2002). Soil fertilization significantly impacts global food production even after the discovery of Haber-Bosch nitrate synthesis in the early 20^th^ century (Erisman et al., 2008). The ability to produce synthetic nitrates made nitrogen fertilizers readily available. Unfortunately, the prolonged or the overuse of inorganic nitrogen fertilizer is unsustainable, disruptive of the N-cycle in ecosystems, and thereby negatively impacting the environment (Kinzig P. Ann, Socolow, 1994; Vitousek et al., 1997). This has led to high advocacy for scientific advances and policy changes which control the environmental impacts of agriculture (Tilman, 2001).

Increasing legume production is a long-term sustainable soil amendment strategy that could further contribute to ameliorating nutritional gaps in protein diet, particularly in low-income economies such as sub-Sahara Africa (Dakora and Keya, 1997; Duranti and Gius, 1997). In sub-Sahara Africa, the tropical savanna landscape is characterized by low soil nitrogen, high ambient temperature, and drought all of which impede crop production. While several resources tend to limit plant growth simultaneously under natural conditions (Chapin et al., 1987), soil fertilization remains the advocated intervention to improve crop yield in savanna agriculture (FAO, 2005; Martey et al., 2013). However, soil fertilization in the absence of government subsidies and international aid programs could further increase the cost burden of farming on low-income farmers. Moreover, a large percentage of inorganic fertilizer leaches down into the subsoil where is not taken up by the crop plants making this expenditure less efficient. Furthermore, fertilizer application does not necessarily address the critical challenges that confront field-grown crops. Although nitrogen fixation offers leguminous plants a competitive advantage in savanna ecosystems, nitrogen (N) fertilization is long known to decrease N-fixing rates in legumes (McAuliffe et al., 1958; Salvagiotti et al., 2008). However, it is unclear whether N-fertilization is the optimal intervention for improving legume mono cropping under savanna climate, given the importance of N-availability in legume development.

The functional relevance of N during legume development underscores the need for investment in securing allocation from pooled sugar resources. Legumes allocate photoassimilates to symbiotic microbes under limited soil N conditions, to increase N-fixation rates. Likewise, during abiotic stress, plants convert photoassimilates into protective metabolites such as compatible solutes (osmoprotectants) to alleviate stress impacts such as the evolution of reactive oxygen species under drought conditions. Consequently, both abiotic stress response and nitrogen needs in savanna legumes may eventually compete from a common carbon resource pool.

These facts underpin the importance of allocation decision in plants (Lerdau and Gershezon, 1997); support the need to re-assess, and test existing theories to ascertain and inform intervention in situations where both nitrogen limitation and abiotic stressors represent the reality in field conditions as pertain in the sub-Saharan Guinea savanna.

According to the Resource Availability Theory (RAT), plant species in low resourced environments invest in defensive allocation (Coley et at. 1985) such as the synthesis of compatible solutes from sugars as a trade-off to growth. On the contrary, the Growth Differentiation Balance Hypothesis, GDBH (Loomis, 1932) is based on a higher physiological sensitivity towards growth such that allocation favors growth. For instance, as in the case of nitrogen limitation where excess photoassimilates are allocated towards plant defense. A bridging or intermediate theory of the two is the Carbon-Nutrient Balance Hypothesis, CNBH (Bryan et al. 1983) which advances that more carbon, for instance, is allocated under nitrogen limitation to both acquire more nitrogen and protect nitrogen resources. While ecological theories encapsulate the behavior of natural ecosystems, agricultural practices or interventions may not, particularly in the case of species self-regeneration and biotic competition. Rather, agriculture remains a major land use threat to natural ecosystems yet, an indispensable basic support of human survival. In essence, agroecology as a sustainable agronomic practice can be consistent with predictable ecological patterns to ascertain predicable outcomes. Although crop production often tends to be monoculture, a monoculture agronomy in itself is comprehensible from an ecological point of view as a single species interacting with each other in disturbed growing space. Agronomic interventions that seek to optimize the crop-environment feedback dynamics towards desirable productive outcomes can implement this perspective. Of particular interest is how legumes partition resource allocation when growth conditions demand multiple needs such as N-fixation and drought stress responses.

Where grain yield is the primary goal of the agronomist, intervention strategies for crop development that regulate responses can utilize knowledge of legume allocation strategies.

N-fixation capacity is an adaptation to higher nitrogen need in legumes and/or serves as a competitive trait over non-leguminous species in N-limited environments. Under limited soil N, increased N-fixation response is expected. Where field conditions necessitate, legumes may employ additional responses to deal with different environmental stresses (Bryan et al., 1996). This may increase demand on pooled carbon resources, possibly resulting in a share or shift in carbon allocation towards more vulnerable or demanding sinks. Deciphering the limiting stress factor under multiple-stress conditions is essential for developing efficient and cost-effective agroecological interventions for stressful environments.

In this paper, we propose the Limiting-Stress-Elimination Hypothesis (LSEH) and test it using non-hormonal biostimulant, varied concentrations of a polyol-based leaf osmoprotectants as exogenous metabolites and nitrogen fertilizer as interventions to cowpea response under Guinea savanna conditions.

### 1.1 Hypothesis

LSEH assumes that plants exposed to multiple physiological stressors will relatively show higher physiological sensitivity and eventually, increased productivity when an intervention eliminates/ameliorates the most limiting stress. This hypothesis stems from Carl Sprengel and Justus von Liebig’s Law of the Minimum which states that growth is controlled by the limiting factor or scarcest resource (Liebig, 1840; van der Ploeg et al., 1999). Agronomic intervention(s) here refers to intended action(s) that remedies condition(s) that impair crop plant productivity. Thus, under multi-stress scenarios where a plant is constrained by more than one stress factor, an intervention that eliminates the most threatening stressor may likely alleviate the stress and maximize productivity when it is economically impractical to eliminate all prevailing stressors.

On the decrement from the maximum hypothesis (Mitscherlich, 1909), maximum productivity requires relieving all decrements from the optimal level for each nutrient, but we can use the term stressor as mineral nutrients are not the only stressors that limit productivity.

Particularly in the Guinea savanna of West Africa, where leguminous crops are predominantly constrained by limited soil nitrogen and osmotic stressors (Fig. 1), the simultaneous treatment to eliminate multiple abiotic stressors may increase farming costs beyond economic practicality.

**Fig 1.**
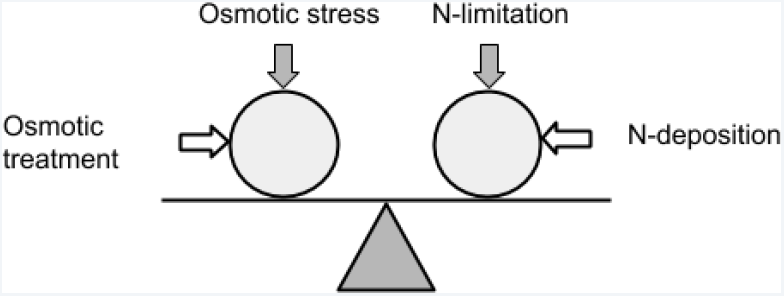
A schematic of the limiting-stress-elimination hypothesis (LSEH). Vertical arrows represent imposed stress (Osmotic stress and N limitation). Horizontal arrows indicate applied intervention (osmoregulation and nitrogen deposition). Circles represent plant organism. Balanced plane implies a notion of significance or pedigree of stress impact. At equilibrium, the stressor impact on productivity is assumed similar. Without any intervention, stress impact is can be assumed synergetic or difficult to detect. However, when defined interventions are simultaneously applied, a shift in slope or tilt distinguishes the relative magnitude of stressor and limiting growth factor.

Although legumes can biologically fix N, they depend on prior invested carbon reserves much as a metabolic response to osmotic stress. The LSEH hypothesizes that, eliminating a relatively more threatening stress among multiple stresses, will increase productivity more than alleviating a relatively less threatening stress. Yet again, there is the question of how the *more* limiting stress is determined under field conditions?

In this study, we determine the limiting stress by testing which stress intervention favors higher productivity in cowpea based on biomass allocation and stress response strategy. Further, we examine how results relate or support existing allocation theories. Then we evaluate the implication of the proposed LSEH on leguminous crop production in economically and environmentally resource challenged sub-Saharan savanna.

## 2.0 Materials and methods

Here we describe how we experimentally tested hypothesis with cowpea in the Guinea savanna agroecological of zone in Ghana, West Africa.

### 2.1 Study area

The Northern region covers 40% of Ghana’s (92,456 sq. mile; latitudes 4-12^0^N) land cover by area. Two major ecological zones characterize the region: the sub-humid to semi-arid Guinea savanna and the arid Sudan savanna zones (Gyasi 2015). The Inter-Tropical Conversion Zone controls rainfall seasons in Ghana with one wet season in the north and two wet seasons in the south. Low, erratic rainfall ranging between 150-250 mm/month in a single dry season to 1100-1200 mm/month in a single wet season characterizes the study area. Mean monthly temperature during the growing season ranges between 26-30 °C (Buah and Mwinkaara 2009). Although the study was conducted at the onset of the growing season, the total rainfall recorded during the study depicts a typical Guinea savanna dry season weather (Table 1).

**Table 1.** Weather records and soil texture composition of experimental site

| Weather |  | Soil composition (%) |  |
| --- | --- | --- | --- |
| Total rainfall (mean) | 203.8 (6.6) mm/day | Sand | 55.8 |
| Average sunshine | 5.8 Hrs | Silt | 41.96 |
| Average mean Temp | 26.7 °C | Loam | 2.2 |
| (Max\|Min) | (29.7\|23.6) °C |  |  |
| Average mean RH<br>(Min\|Max) | 84%<br>(93\|75)% |  |  |

Beneath the surface soil is a shallow depth of cemented layer of iron pan and savanna ochrosols. The soil is humus-deficient due to low organic matter content from sparse vegetation. Iron pan impedes water penetration causing water logging in the peak rainy season and complete drying in the dry season. Hence, soil nutrient availability is a major constraint for agricultural production (Runge-Metzger and Diehl 1993; Gyasi 1995).

### 2.2 Preparation of field, plants and treatments

Three experimental blocks, each containing four distinct raised soil beds of dimension 2m × 8m each and interspaced by a meter gap were prepared across a slope in a series-like fashion. Each bed in a block represented a treatment group based on a Randomized Complete Block Design.

Each bed was seeded with cowpea (IT97K-499-35, Songotra) and later thinned to 40 plants per bed after germination at 80cm × 40cm spacing. Treatments were assigned randomly within and between blocks. Given the conditions of Guinea savanna, we will use the term osmotic stress to represent both drought and heat stress due to their occurrence duality (Dwivedi et al., 2018) or specifically to represent conditions such as low soil moisture, atmospheric vapor pressure deficit and salinity for their similarity in physiological responses (Chaves et al, 2009; Munns 2002). The first osmotic treatment constituted two different concentrations; 0.25g/L (HH25) and 0.5 g/L (HH50) of the osmolytic polyol, Hexane 1, 2,3,4,5, 6-Hexol (C_6_H_14_O_6_) of molecular weight 182.17 g/mol obtained from Sigma Alderich Co. These represent separate treatment groups for statistical convenience. In addition, a non-hormonal biostimulant (Biost.) comprising as osmotic stress treatment, made up of; 2-Amino-5-guanidinopentanoic acid, L-ascorbic acid, thiamine (3-[(4-Amino-2-methyl-5-pyrimidinyl) methyl]-5-(2-hydroxyethyl)-4-methyl-1,3-thiazol-3-ium nitrate) and *cis*-1, 2, 3, 5*-trans*-4, 6-cyclohexanehexol (*myoi*nositol). The formulation was developed separately in the lab based on several allocation and yield trials in glasshouse experiments on allocation study model plant, *Raphinus sativus*.

Further, 5 g/L nitrogen was prepared from 98% ammonium nitrate (NH_4_NO_3_) as inorganic nitrogen treatment (N Fertiliser). All controls groups were treated with distilled water. Each plant replicate received 50 cm^3^ of treatment dose biweekly until the onset of pod formation. Nitrogen was applied directly to the soil around the stem of plants while *HH25, HH50, Biost*. in addition, control were each applied to both foliage and soil.

### 2.3 Plant sampling and measurement

To minimize the influence of growth defects, 15 best germinates out of 40 ranging ∼9-11cm in height were tagged after establishment. Leaf measurements were conducted during active photosynthetic period in the morning (PPDF ranged from 900-1200 m molm^-2^s^-1^). The Leaf temperature, chlorophyll fluorescence and chlorophyll concentration of the youngest fully expanded leaf of each tagged plant was measured two weeks after treatment during the vegetative, flowering and podding growth phases of cowpea.

#### Biomass and Nitrogen-fixing trait

Whole plants were carefully removed from soil bed, the soil debris gently brushed off. Root nodules per plant were carefully removed, counted, weighed and re-weighed after oven drying to a constant weight at 60 °C to obtain total weight of nodule per plant and total number of nodules formed per plant. Means for each treatment groups are reported in section 3. Furthermore, whole plants with intact pods were oven dried to constant weight to obtain total biomass, then followed by separate measurement of pods.

#### Leaf temperature

Leaf temperature, *T*_*l*_was measured as an indicator of plant stress (Carroll et al., 2017; Rodríguez et al., 2015; Udompetaikul et al., 2011). Variations in moisture stresses are known to significantly alter leaf temperature, causing it to deviate from ambient temperature (Wiegand and Namken, 1966). *T*_*l*_ was measured by a laser guided infrared thermometer (ST60 ProPlus™ Raytec, USA) with 30:1 optical resolution, −3°C-600 °C temperature range, 8-12μm spectral responses at 500 msec, and 0.10-1.0 emissivity. The device collects and focus energy optically emitted, reflected, and transmitted by the leaf surface on a detector, then converts to ambient temperature readings in °C with 0.07 °C error margin.

#### Chlorophyll fluorescence

The stress manifested in plants can be determined by observing the leaf photosystem II (PSII), which is a protein complex in the light dependent reactions of plant photosynthesis. PSII damage is the first manifestation of stress in plant leaves (Maxwell and Johnson, 2000) and the efficiency of this state is referred as photochemical efficiency. Maximum quantum yield at PII is determined based on the fluorescence ratio, *F*_*v*_ *F*_*m*_,^−1^ a well-studied phenomenon in plant physiology (Rodríguez et al., 2015) and well established physiological indicator of plant tolerance to environmental stress including drought and heat (Dwivedi et al., 2018; Maxwell and Johnson, 2000). Plant stress that affects the photosystem II is determined based on a calculated variable fluorescence (*F*_*v*_) to maximal fluorescence (*F*_*m*_) ratio, using a chlorophyll fluorescence device (model OS-30p, Opti-Sciences, Inc.). To measure 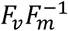, leaf samples were dark-adapted by placing leaves inside plastic cuvettes that prevented light incidence on the leaf for 30 minutes to excite pre-photosynthetic antenna by a weak modulated light. At this stage, photosystems II are maximally oxidized and the minimal fluorescence, *F*_*0*_, was measured. On light saturation at maximum light exposure, the maximum fluorescence, *F*_*m*_, was determined and 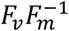 calculated as:

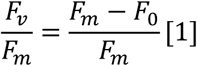

Where *F*_*v*_=*F*_*m*_–*F*_0_ is the variable fluorescence.

#### Chlorophyll content

Leaf chlorophyll content is a physiological trait associated with drought and heat stress (Dwivedi et al., 2018). A handheld chlorophyll-detecting device, SPAD-502 (Konica Minolta Inc.), measured leaf chlorophyll. Leaf was placed in between an attached in between a receptor window and a measuring head on which is pressed to measure chlorophyll at 650 and 940nm in SPAD units at ±1.0 accuracy (see Richardson, et al, 2002).

### 2.4 Statistical assumption

The null hypothesis (H_o_), no differences in population means:*T*_1_ = *T*_2_= *T*_3_= *T*_4_; P = 0.05.

The null hypothesis (H_o_), no differences in population means: *T*_1_= *T*_2_= *T*_3_= *T*_4_; P < 0.05, where *T*_*n*_indicates treatment type *n*.

### 2.5 Statistical analysis

The Lme4 package in R (Bates et al., 2015) was used to perform Mixed Effect Modelling (MEM) by restricted maximum likelihood t-tests. MEM performs ANOVA on repeatedly measured variables and estimates fixed and random sources of variation. To know where significant differences occurred, we tested general linear hypotheses by multiple pairwise comparisons of means based on Tukey HSD tests on normally distributed data. All statistical analyses were conducted using the R programming language and environment (R Core Team).

## 3.0 Results

### 3.1 Effect of treatments on biomass, podding capacity and root nodulation

Mean biomass of treated cowpea was significantly different (P<0.001, Table 2). Non-hormonal biostimulant had highest biomass output, 45% followed by HH25 (17%), HH50 (13.3%), N Fertilizer (13.1%) and controls. Nodulation capacity, determined by means of the total number of nodules formed per cowpea was significantly different (P < 0.01) among treatment groups. N treated cowpea relatively formed the highest number of nodules per root as well as the highest nodule weight (Fig 2, c & d). Thus, we reject the null hypothesis that there is no difference in the mean effect of administered treatment interventions on cowpea growth with respect to biomass and nodulation capacity.

**Table 2.**
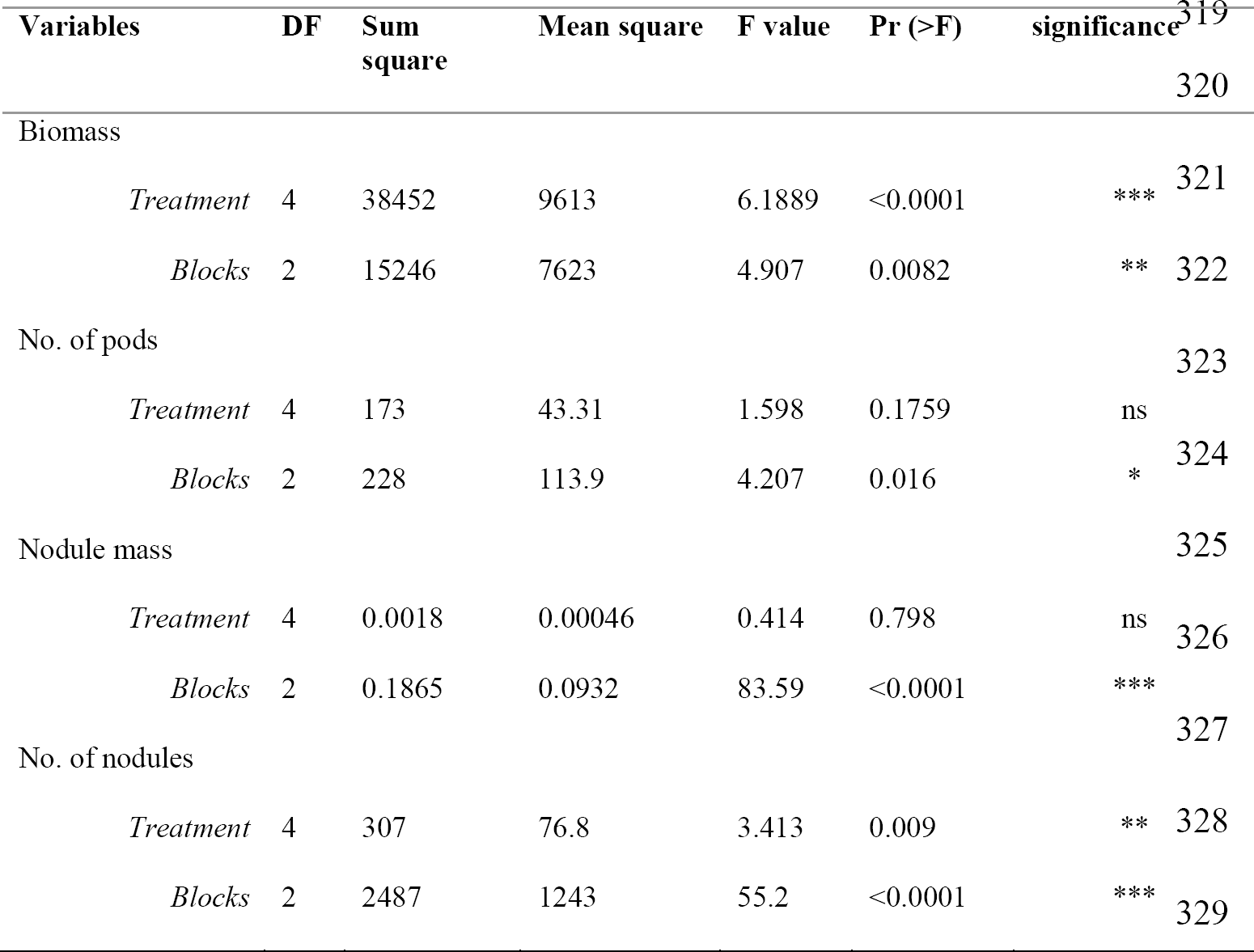
Summary ANOVA of mean response of treatment groups.

**Fig 2.**
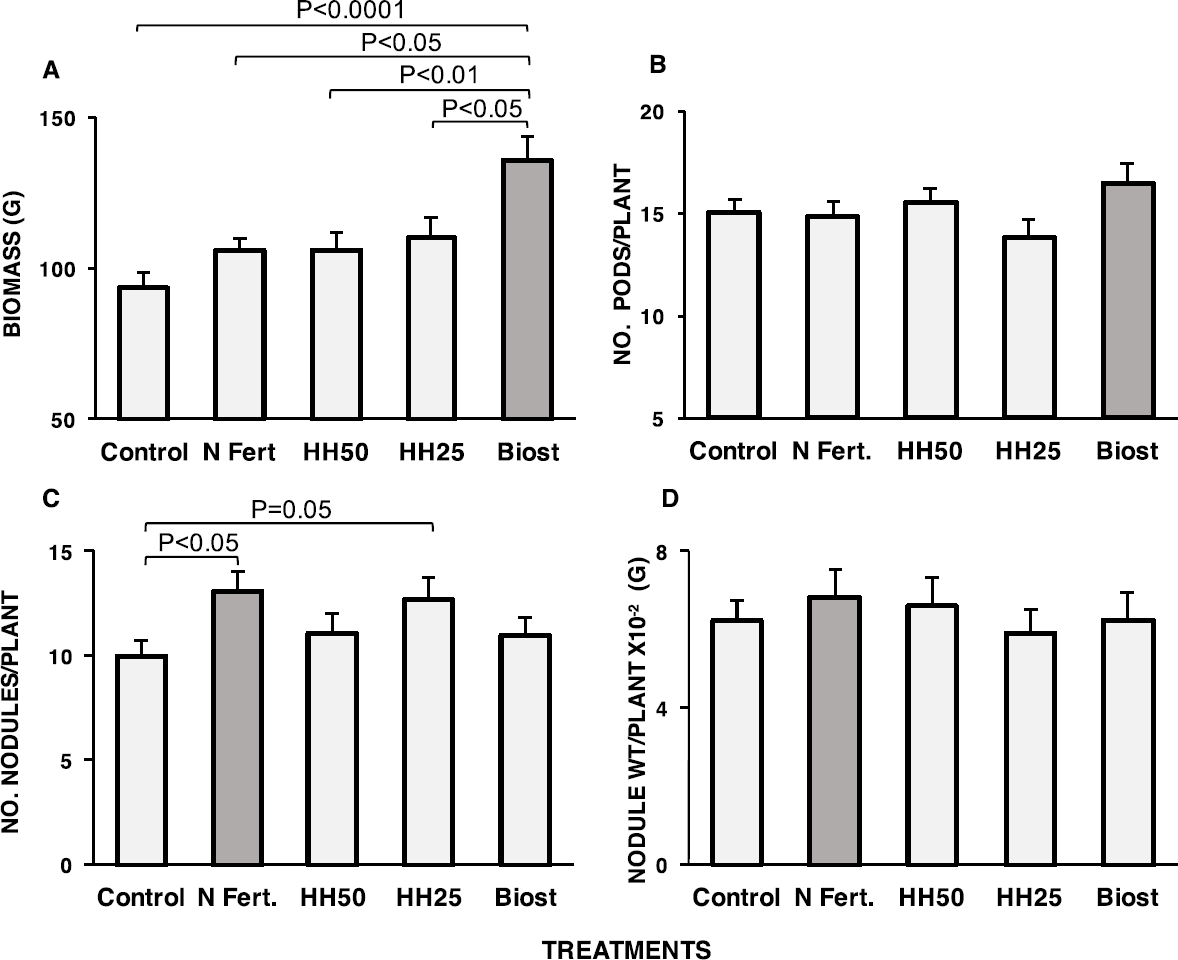
Means of total biomass, grams/plant (A), mean number of pods per plant (B), mean number of root nodules formed by roots per plant (C) and mean weight of root nodule per plant (D) in response to cowpea treatments; control, N Fertilizer (N Fert), sorbitol (HH25 & HH50) and non-hormonal Biostimulant (Biost). P-values on bars indicate Tukey HSD significance between paired groups.

The treatment effects on both the mean number of pods formed per cowpea and mean nodule mass per cowpea root nodules were not significant (P>0.05, Table 2). However, the shows non-hormonal biostimulant treatment relatively yielded more pods per plant.

### 3.2 Effect of treatments on leaf physiological responses

The mean effect of treatments on leaf chlorophyll content in cowpea did not differ significantly, however, the difference (P=0.05) in leaf chlorophyll responses to HH25 and HH50 indicates that the concentration of exogenous leaf osmoprotectants is important to chlorophyll response in savanna cowpea. Maximum quantum yield, 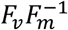, in response to N differed significantly from osmotic treatments and controls (Table 3). Leaf temperature, *T*_*l*_significantly declined in N fertilized cowpea relative to controls and osmotic stress treatment groups (Fig 3c, Table 3).

**Table 3.** Significant pairs from multiple mean comparisons based on Tukey HSD test from a linear Mixed Effect model fit. Except reported, all other possible mean-pairs were not significant.

| Parameter variables | Estimate | Std error | z value | Pr(>\|z\|) | significance |
| --- | --- | --- | --- | --- | --- |
| FvFm<sup>-1</sup> |  |  |  |  |  |
| Fertilizer-Biost | 0.0368 | 0.0128 | 2.859 | 0.033 | * |
| Fertilizer-HH25 | -0.0333 | 0.0113 | -2.926 | 0.028 | * |
| Fertilizer-HH50 | -0.0399 | 0.0113 | -3.506 | 0.004 | ** |
| Chlorophyll content |  |  |  |  |  |
| HH25-HH50 | -2.5979 | 1.0300 | -2.522 | 0.05 | . |
| Leaf temperature, $T_l$ | | | | | |
| *Fertilizer-Biost* | -1.1436 | 0.2867 | -3.989 | <0.001 | *** |
| *Fertilizer-control* | -1.3616 | 0.2554 | -5.345 | <0.001 | *** |
| *Fertilizer-HH25* | 0.7694 | 0.2548 | 3.020 | 0.021 | * |
| *Fertilizer-HH50* | 0.9004 | 0.2548 | 3.534 | 0.003 | *** |

**Fig 3.**
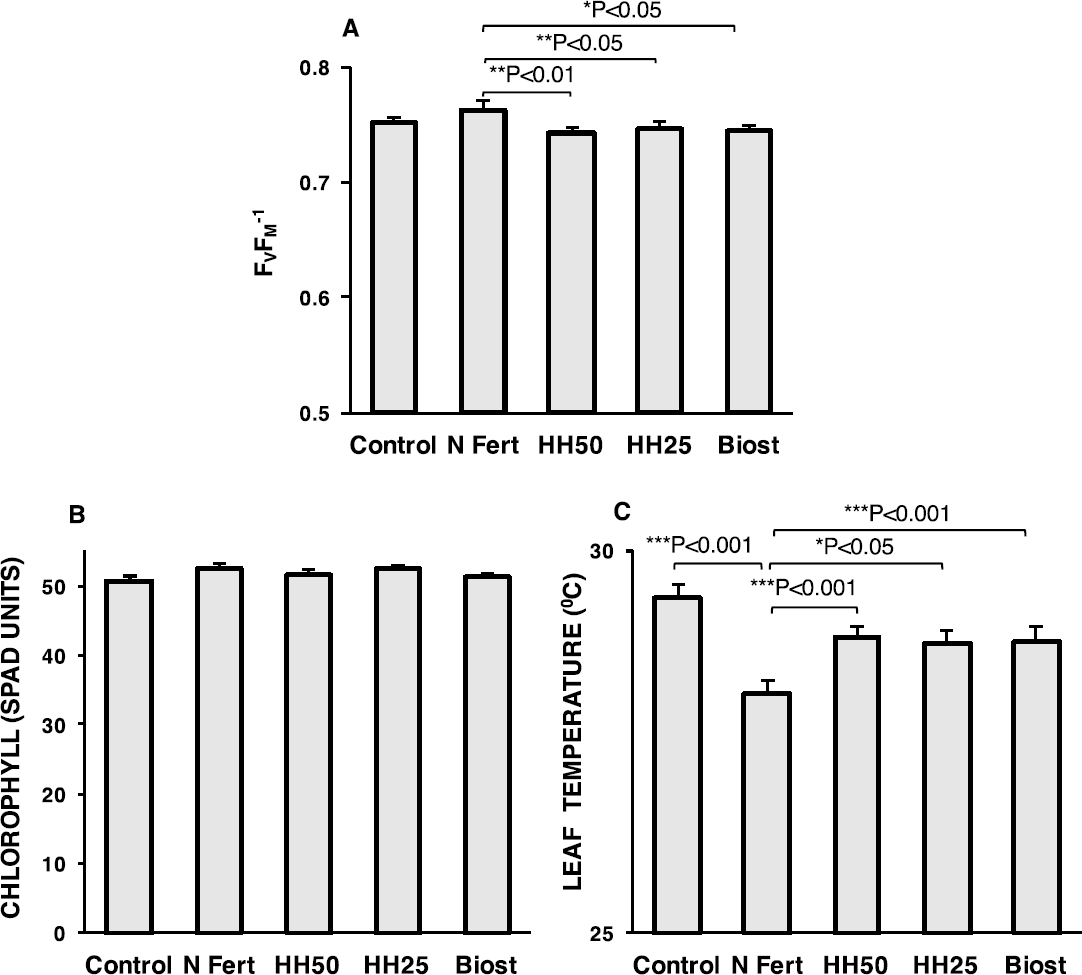
Leaf physiological response of cowpea to osmotic treatments (HH25, HH50 and Biost) and N treatment (N Fert.). Bar graphs represents total oven dry biomass (A), leaf chlorophyll content (B), chlorophyll fluorescence ratio (C) and leaf temperature (D).

For evaluating the effect of cowpea treatment with respect to stress response, we show the underlining relationship between selected indicators from our study data (Fig 4).

**Fig 4.**
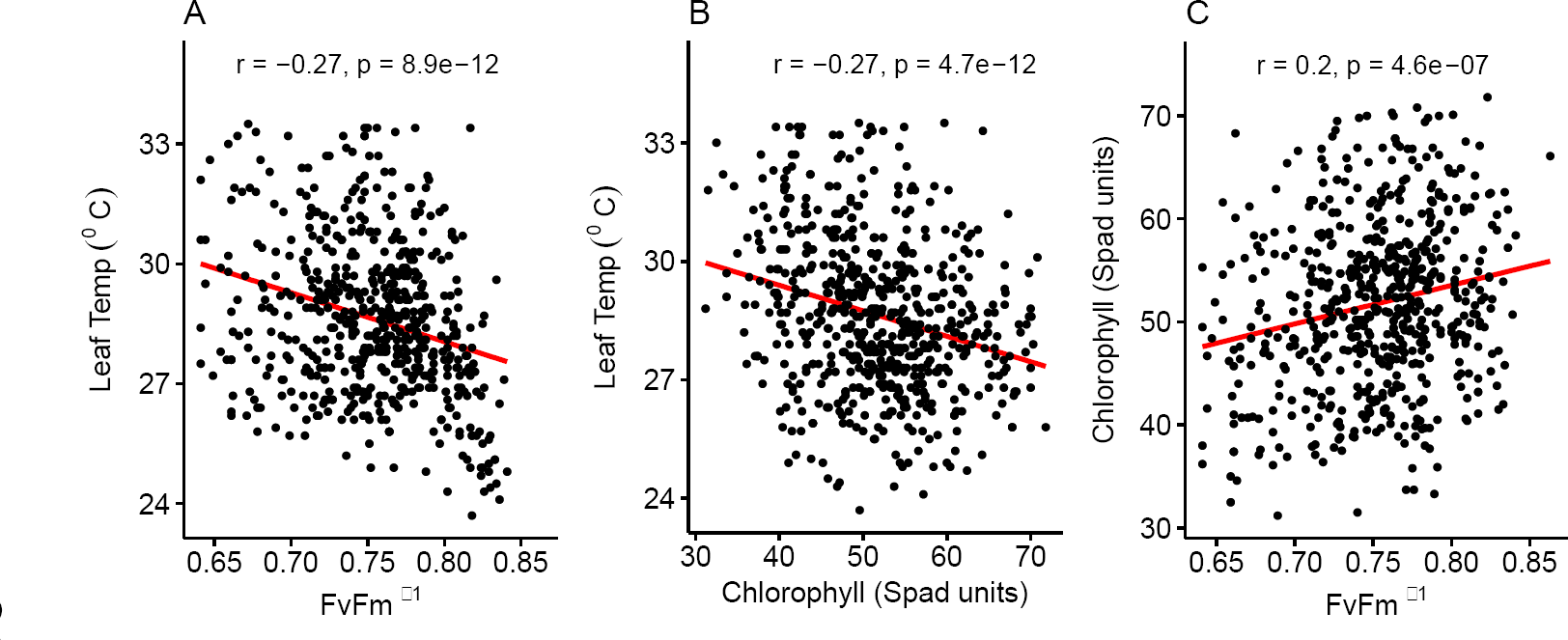
Correlation of stress indicator variables. A) Leaf temperature (Leaf Temp °C) and 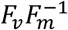. B) Leaf temperature (Leaf Temp °C) and Chlorophyll content. C) Chlorophyll content and 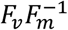.

## 4.0 Discussion

### 4.1 Biomass and physiological responses of cowpea to osmotic and N treatments

Cowpeas cultivated in the savanna are exposed to multiple environmental stressors. Environmental stressors in the context of the plant-environment-feedback nexus, shape the fundamental developmental trajectory and productive success of plants. In cowpea production, low soil fertility and drought constitutes abiotic factors that constrains production (Roberts, 2013). Although legumes are adapted to low N conditions, N-fixation comes at a cost. Under savanna conditions, other competing abiotic factors may inevitably constrain biological N-fixing capacity of nodulating legumes, and consequently limit their productivity. Hence, we hypothesized that where cowpea is constrained by multiple factors, elimination of the relatively more limiting factor will result in relative increased productivity. We measured productivity by total biomass output, complemented by pod yield. Given relatively higher biomass, pod yield and less investment in root nodules in osmotic treated cowpea, we conjecture that osmotic stress is more limiting than N-limitation in Guinea savanna cowpea.

In this study, a healthy physiological status is attributable to higher chlorophyll content, 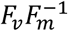 and lower leaf temperature (Fig 4).

While we expected higher N fixing capacity and 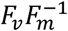 to increase productivity, the response was relatively lower in nitrogen treatment than osmotic stress treatments. The relatively lower biomass and increased nodulation response to N suggests a trade-off for increased investment in the need for physiological response and N-fixation (Fig 2c, 2d). Contrary, the relatively higher biomass and leaf stress responses to osmotic treatments support the conjecture that exogenous stress compounds alleviate the need and the cost of metabolic response, thereby directing more allocation towards biomass output.

### 4.2 Effect of non-hormonal biostimulant and sorbitol on cowpea

Biostimulants, since its invention, definition and development over the past four decade have been shown to improve stress resistance and promote plant development (Russo and Berlyn, 1991). Although drought avoidance by hydraulic controls (Hall and Schulze, 1980; Turk and Hall, 1980) and paraheliotropism (Schakel and Hall, 1979)(also observed in this study) persist in cowpea, osmoadaptation through metabolite accumulation has been reported recently as a more conservative strategy in cowpea (Goufo et al., 2017). From our results, we conjecture that exogenous osmotic stress compounds may alleviates the metabolic need and cost towards osmoadaptation, thereby making available more photoassimilates for biomass allocation.

Among 88 metabolites studied in cowpea cultivars by Goufo et al. (2017), only proline, galactinol and a quercetin derivatives were reported beneficial to yield, whereas *myo*inositol and arginine were not. Contrary from our findings, *myo*inositol and arginine, both key components of the non-hormonal Biostimulant to impacted cowpea yield. Further, we report the first yield effects of exogenous sorbitol on cowpea. While sorbitol does not constitute reported osmolytic compound in osmotic stressed cowpea, its yield effect in our study suggests metabolite-yield relation may be non-specific.

### 4.3 Implication of the Limiting-Stress-Elimination-Hypothesis in Savanna legume agroecosystems

According to the proposed LSEH, when the limiting stress is eliminated, stress impact is relieved (Fig 5), until the next available stress becomes most limiting. The scope of this study here is not to establish the most limiting stress in cowpea. Instead, we have demonstrated which, among osmotic stress and low soil N limits cowpea productivity by showing whose ‘elimination’ (by treatment) relatively improve productivity.

**Figure 5.**
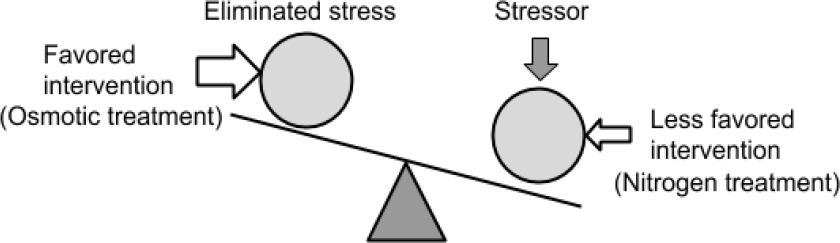
A schematic of limiting-stress-elimination hypothesis (LSEH). Here, a favored intervention shows sensitivity to stress response by plant, eliminate or alleviates the impact of stressor, and results in increased productivity indicated by upward tilt analogous to a “lighter weight effect”.

Agroecology is a sustainable approach to crop production by additionally promoting the ecological footprint of agronomy. Increasing legume production constitute a long-term food security initiative and ecological restoration strategy whereby legumes improve soil N, increase availability of percent arable lands and the possible recruitment or cultivation of less competitive but essential non-legume plants or crop species in the savanna. In our study, we measured N-fixation marker by nodulation capacity and mean nodule dry mass. While N treatment relatively increased nodulation capacity significantly, the effect of N treatment and osmotic treatment on mean nodule dry mass did vary significantly. Thus, exogenous osmotic stress treatment constitute a promising alternative for increasing cowpea production without impairing their biological N-fixing capacity. N-fixation is an important ecological process and more critical, in low vegetated semi-arid ecosystems such as Guinea savanna. However, the necessity to use N fertilizers as supplemental input to maximize legume yield should depend on whether additional inorganic N supplement addresses the prevailing limitation to growth. Evidence exist to question the relevance of N fertilization in cowpea. For instance in a study, bacteria inoculated cowpea more than N-fertilized cowpea improved grain productivity by increasing biological N-fixation (Martins et al. (2003). In this instance, regardless of N supplement, increased biological N-fixation through bacterial inoculation accounted for higher relative growth. Essentially, the metabolic cost of microbial aided N-fixation may be lower than the metabolic cost of converting inorganic N fertilizers to chemically useful forms impact higher crop yield. Similarly, in our study, minimizing the cost of stress allocation may have influenced yield. Thus, we conjecture two possibilities that accounts for increased growth in cowpea through exogenous stress treatments. First, osmotic treatments alleviate the metabolic need therefore the cost for osmotic stress allocation, resulting in more readily available carbon reserve. Second is the consequential decline for the need to share from the carbon pool, photosynthate allocation to neutralize stressors such as the synthesis of compatible solutes towards osmoadaptation. In both conjectures, more carbon is made ready for biomass allocation than towards defense.

In allocation theories, plants constrained under limited resources broadly tend to exhibit either defensive or reproductive allocation strategy. LSEH is an intervention for defensive allocation that satisfies the Resource Allocation Theory. Contrary, N fertilization represents an ideal intervention where plant strategy satisfies the Carbon-Nitrogen-Balance hypothesis. The LSEH differ from the Spregel-Liebig concept only by the intervention approach whereby in the latter, supplementing nutrient deficient plants by sufficient amounts of various limiting nutrients may have been the considerable intervention. However, as exception to legumes with innate capacity to fix a degree of atmospheric N_2_ even when soil N is limited, the LSEH focuses on the principal limiting stressor, in this case, osmotic stress. It can be inferred from biomass responses that a less productive outcome may ensue when a relatively more limiting factor persist while an intervention is targeted at a relatively less limiting factor, such as soil N fertilization, as the common practice by cowpea growers in Northern Ghana. Hence, it becomes imperative for the agronomist to determine what constitute the measurable limiting factor to any crop of interest. When that is determined, management techniques and tools that improve resource use efficiency become the next relevant agronomic asset (Chaves te al, 2009). The LSEH utilizes a cost-benefit utility that should inform decisions and potential innovations entrenched in the principle of efficiency and sustainability in agricultural practices particularly, in regions challenged by both limited economic and environmental constraints.

We have demonstrated through our study that intervention towards osmotic stress alleviation can improved cowpea biomass allocation compared to N fertilization. In the search for sustainable agroecological solutions, LSEH can assist agronomist to decipher appropriate interventions optimal for improving productivity.

## 5.0 Conclusion

Crops growing in Guinea savanna are constrained by a myriad of abiotic stressors such as soil N-limitation and osmotic stressors such as drought and high temperature. Eliminating such constraints may ideally require multiple interventions. However, limited economic incentives makes this impractical, especially in low-income economies. Fertilizer remains the predominant approach to maximize crop productivity yet can be economically and environmentally unsustainable. This study proposes and tests the Limiting-stress-elimination hypothesis (LSEH), which posits that eliminating the limiting factor sufficiently maximizes productivity where other competing constraints exists.

In our study, we treated cowpea with a non-hormonal biostimulant and exogenous metabolite to alleviate osmotic stress. Further, we supplemented cowpea with inorganic N to alleviate soil N-limitation. From the results, osmotic stress treatments more than N treatment increased cowpea productivity, assessed by total biomass output and pod yield.

The study further implies that osmotic stress is more limiting than N in cowpeas under Guinea savanna conditions. The study validates LSEH, which underscores the importance crop nutritional intervetion that targets non-nutrient limiting abiotic stressors in-leu of fertilizer inputs even for N-demanding systems such legumes.

